# Agonistic behavior and fear response during consecutive encounters in female fish

**DOI:** 10.1101/2025.02.24.639906

**Authors:** Chiara Salustri, Luciano Cavallino, Mariano Brasca, Andrea Pozzi, María Florencia Scaia

## Abstract

While consecutive agonistic encounters modulating aggressive behavior are well studied in males, the extent to which social behavior can be altered by fear sensing during consecutive agonistic encounters remains unexplored. A well-established paradigm to assess fear sensing is the exposure to alarm substances (AS) from conspecifics in Ostariophysi fish, such as zebrafish, which causes several physiological and behavioral changes associated to distress. Considering that the exposure to AS induces changes in individual behavior associated to fear response, the aim of the present work is to examine whether repeated exposure to AS alters both agonistic behavior and fear response in female zebrafish. Moreover, since female aggression is understudied when compared to males, this study also assesses possible differences according to the reproductive stage. We performed intrasexual dyadic encounters between female zebrafish to determine if agonistic behavior is altered during two consecutive contests and by the presence of AS. When comparing agonistic behavior in different reproductive stages, results suggest there are no differences in latency and in freezing. Regarding time of aggression, while there are no differences between contests in prespawning or postspawning, significant differences are detected between postspawning dyads and mixed dyads with both females in different reproductive stages. Results suggest that exposure to AS reduces female motivation to engage in an agonistic encounter while aggressive behavior is still maintained despite sensing AS as potential threat, regardless of corresponding to the first or second contest. Finally, similar to agonistic behavior, the effect of fear sensing on individual behavioral parameters such as distance, mean velocity and freezing is also observed during a first and also a second exposure to AS. To the best of our knowledge, this is the first evidence assessing how consecutive agonistic encounters with a real opponent can be altered by repeated exposure to AS.

**Summary statement:** We assess whether repeated exposure to distress alters both agonistic behavior and fear response in female zebrafish, by performing consecutive agonistic encounters with a real opponent exposed to alarm substances.

## 1. Introduction

Individuals living in social hierarchies engage in agonistic encounters through aggressive behavior, which induces experience-dependent shifts in social status. Social behavioral outcomes integrate not only different internal physiological cues, such as hormonal and neuroendocrine pathways, but also experience-related memories and the sustained sensing of a complex constantly changing environment (Pandolfi and Silva, 2021, Filby et al, 2010, Maruska et al., 2022, Quintana et al., 2021, Kareklas et al., 2023). In this context, consecutive agonistic encounters in *Danio rerio* (zebrafish) males modulate the aggressive behavior and suggests a formation of a long-term social memory, related to recognizing a particular opponent and/or features of previous fights (Cavallino et al., 2020, 2024). However, the extent to which social behavior can be altered by fear sensing and distress during consecutive agonistic encounters remains unexplored in both sexes.

A well-established paradigm to assess different aspects related to fear sensing is the exposure to alarm substances (AS) from conspecifics in fish (Jesuthasan and Mathuru, 2008). Fear response in Ostariophysi fish, such as zebrafish, can be induced by the alarm substances produced by epidermal club cells (Smith, 1992, Pfeiffer, 1977). Club cells are believed to exert multiple functions (Zaccone et al., 1994, 2001, Chia et al., 2019) and, in this sense, when fish are injured, they passively release alarm substances from their skin, inducing a fear response in conspecifics that consists of erratic movements and freezing behavior (Speedie and Gerlai, 2008). This fear response induces an increase in cortisol levels and stress (Egan et al., 2009), thus suggesting that the exposure to AS is a very appropriate paradigm to study distress caused by fear sensing. Since the AS is a known fear-eliciting stimulus in fish, it has been extensively used to study different manifestations of social information use in threat perception, such as social buffering, social transmission (contagion) and facilitation (Oliveira and Faustino, 2017). Interestingly, evidence in zebrafish suggests a shared evolutionary origin for social buffering of fear in vertebrates, which is the reduction of fear response in presence of conspecifics (Faustino et al., 2017), and a shared evolutionary conserved role for oxytocin as a regulator of emotional contagion or empathy (Akinrinade et al., 2023a). A recent study has identified two compounds in the zebrafish skin extract, ostariopterin and daniol sulfate, acting as “danger” and “conspecific” signals respectively (Masuda et al., 2024). These compounds activate distinct glomeruli in the olfactory bulbs, and they induce robust alarm reaction in zebrafish only when they are concomitantly applied together. Although exposure to alarm substances has been increasingly used to study various aspects of social behavior related to threat perception, it is unclear how sensing potential danger may affect agonistic behavior in subsequent encounters.

The historical association between aggression and males may explain why social behavior during consecutive agonistic encounters in fish has been studied in males but not in female fish yet. Interestingly, although females from different species also display aggressive behavior, female aggression is usually understudied (Borg et al., 2012; Davis and Marler, 2003; Langmore et al., 2002; Renn et al., 2012; Scaia et al., 2020, Pandolfi et al 2021). Information on how aggression fluctuates during the female reproductive cycle in fish is also scarce and varies according to the ethological context. When comparing females in prespawning and postspawning, in the Neotropical cichlid *Cichlasoma dimerus* evidence suggests that prespawning females are more aggressive than postspawning females with eggs or with larvae in a resident-intruder paradigm probably due to the fact that they show higher levels of androgens and estrogens, while in dyadic encounters in neutral aquaria there are no significant correlations between gonadosomatic index and individual aggressive and submissive displays (Tubert et al., 2012, Scaia et al., 2018a, 2023). Since zebrafish is a well-established alternative model to study the neural basis of social behavior, deepening research on female aggression and sex differences in social behavior is the logical next step. In this species females can retain eggs (prespawning) until they are stimulated by a male during courtship (postspawning), and egg-lay can occur repeatedly throughout the same courtship episode (Darrow and Harris, 2004). Even if there are sex-differences in aggression and in the neuronal architecture of intrasexual aggression, both sexes show the same behavioral displays and phases during agonistic encounters (Scaia et al., 2022). Regarding possible differences between reproductive status, since prominent ventral area in prespawning females is usually a very useful characteristic for sexing individuals, evidence on female zebrafish tends to refer to this reproductive stage, thus overlooking variability throughout the reproductive cycle.

Considering that the effect of distress caused by fear sensing has been assessed in individual behavior, the aim of the present work is to examine whether repeated exposure to fear sensing alters both agonistic and individual behavior in consecutive agonistic encounters. For this purpose, we performed intrasexual dyadic encounters between prespawning or postspawning females of zebrafish to assess if agonistic behavior is altered during two consecutive contests and by the presence of alarm substances.

## 2. Materials and Methods

### 2.1. Animals and holding conditions

Zebrafish adults were obtained in a commercial aquarium and they were acclimated in mixed tanks during two months in the Fish Facility in the Facultad de Ciencias Exactas y Naturales (Universidad de Buenos Aires, Argentina). Fish were housed in social tanks of dechlorinated filtered water (1 fish per liter, 20L) at 25 ± 2 ◦C, pH= 7.5/7.8, with a photoperiod of 14 h light (8am-10pm) : 10 darkness (10pm-8am). Fish were fed twice a day with commercial food (Tetra®), except on experimental days. During experimental procedures, female fish (n=94) were maintained in the same water, temperature and photoperiod conditions as in the acclimation period, and two hours after the test they were fed only once with the same amount of food. Holding conditions and experimental design were performed in accordance with international standards on animal welfare (National Research Council, 2011), minimizing any pain and/or discomfort of the animals. All procedures were compliant with the Guide for Care and Use of Laboratory Animals (National Research Council, 2011) and with institutional (*Comisión Institucional para el Cuidado y Uso de Animales de Laboratorio, Facultad de Ciencias Exactas y Naturales, Universidad de Buenos Aires*, Protocol # 75b) and national (*Comit*é *Nacional de Ética en la Ciencia y la Tecnología*) regulations.

### 2.2. Experimental protocol

Fish sex was identified by their coloration and body shape: females are more ventrally rounded than males and display a silver coloration, while males show a yellow/golden coloration, especially in the anal and caudal fin (Paull et al. 2010; Yossa et al. 2013). A total of 47 female-female dyadic agonistic encounters were analyzed for this study. A behavioral paradigm was followed, which has been previously used to study agonistic interactions in male and female zebrafish (Figure 1A, Oliveira et al., 2011; Scaia et al., 2022). For each trial, two size-matched females (with variation in standard length less than 10%) belonging to different social tanks were simultaneously isolated, thus excluding possible previous social interactions between them. No significant differences were found in standard length, total body weight and gonadosomatic index between opponents (Table S1). Animals were paired in size-matched dyads and placed in an experimental arena (3L, 20×12.5×14.5 cm; length x width x height), which was divided into two compartments by a removable opaque partition. Thus, each pair was isolated in an aquaria and both opponents were maintained in visual and not chemical isolation until the next day.

**Figure 1.**
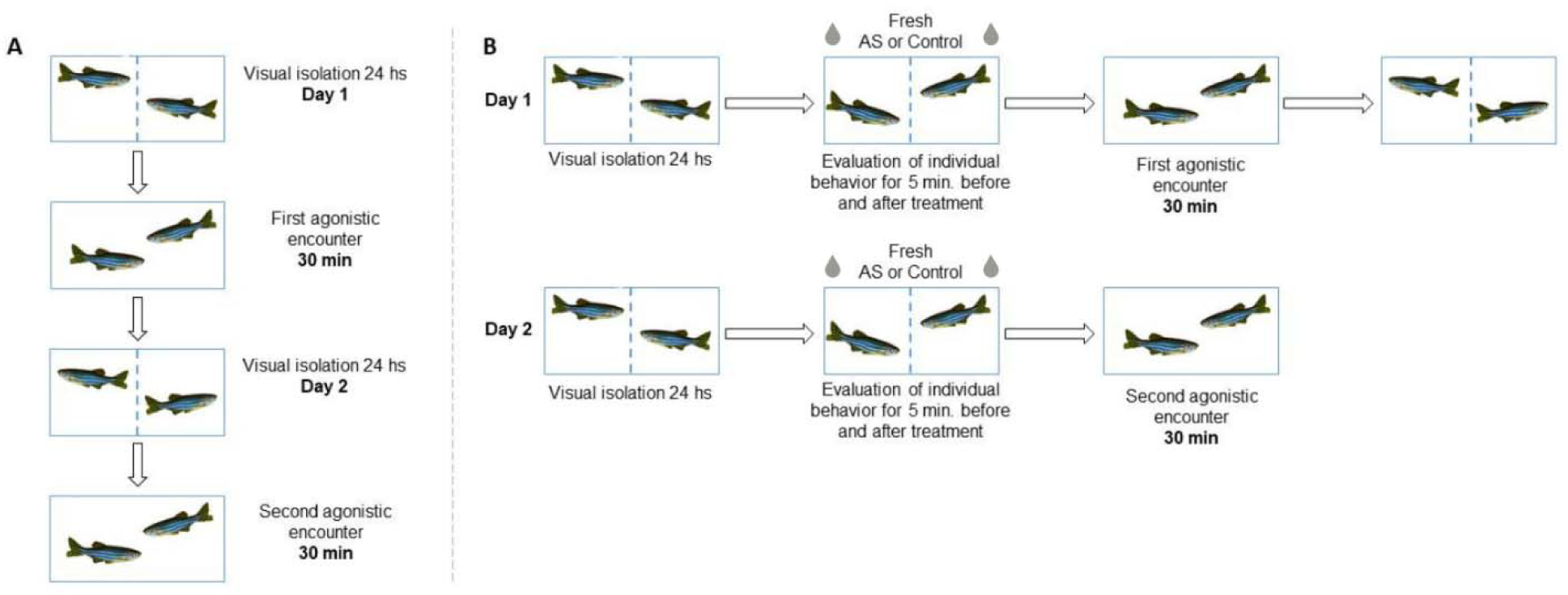
Schematic illustration of experimental protocol for zebrafish fighting. A. After overnight physical and visual isolation, female dyads perform agonistic behaviors for 30 minute. After this first agonistic encounter, opponents are isolated again until the second agonistic encounter on the following day. B. The effect of fear sensing on agonistic behavior was assessed by exposing both opponents to alarm substances (AS) five minutes before starting the first and the second agonistic encounter. Individual behavior was assessed during 5 minutes before and after applying the treatment. To exclude possible effects of volume influx, controls involve only water addition with no alarm substances.

On the next day, the first agonistic encounter was started by removing the partition and allowing both fish to interact for 30 minutes (Teles et al., 2016). The beginning of aggressive behavior occurs when the first aggressive display is observed. As already described in previous studies, female zebrafish typically adopt the same aggressive displays as males, mainly displays, circles, bites, chases and strikes (Scaia et al., 2022). All experiments were performed at the same time of the day (2pm – 4.30 pm) to avoid possible circadian variations. To evaluate repeated agonistic encounters, a behavioral paradigm already used to assess the effect of previous fighting in zebrafish males was followed (Cavallino et al., 2020, 2024). Briefly, once the first trial was finished fish were separated for 24hs with the removable opaque partition. On the next day, the second agonistic encounter was initiated, and both opponents were allowed to interact again for 30 minutes (Fig. 1A).

### 2.3. Alarm substances sampling and treatment

To assess the effect of the distress caused by fear sensing, fish were exposed alarm substances during five minutes before initiating the first and the second encounter. Also, in order to determine whether this effect of fear response on behavior is due to the alarm substances *per se* or to the treatment influx, individual behavior was compared between before and after the stimuli or its control, before performing the agonistic encounter. For this reason, fresh alarm substances were directly sampled before starting the behavioral trials each day. Alarm substances were obtained following the protocol already established and used in this species (Speedie and Gerlai, 2008, Akinrinade et al., 2023a). Briefly, donor prespawning females were euthanized via rapid chilling and placed in a petri dish where fifteen shallow cuts were made on each side of the trunk and washed with 50ml of distilled water. Blood was made certain not to contaminate the solution. Alarm substances from each donor were used for all daily agonistic encounters to control individual variations in donors (e.g. three encounters per day). The alarm substances were kept on ice to avoid degradation during the trials and, thus, distilled water for the vehicle control was kept in the same conditions. While some dyadic encounters received no treatment volume at all (n=17), others were treated with alarm substances (n=15) or with water influx with no alarm substances (n=15, Fig. 1). Treatments (2.5 ml per fish) were administered via a flexible and transparent PVC tubing positioned 0.5 cm above the water level at each half of the tank, five minutes before initiating the agonistic encounters.

### 2.4. Behavioral analysis

The effect of repeated exposure to alarm substances on behavior was studied by assessing individual behavior and also agonistic behavior. To evaluate whether there is an effect of alarm substances on individual behavior, both opponents were maintained isolated in each compartment before agonistic encounters and they were recorded for five minutes before and after applying each treatment in consecutive days (Fig1B). Different parameters on individual behavior were quantified using the commercially available tracking software Ethovision XT© 11.0 (Noldus Inc., The Netherlands). To evaluate agonistic behavior, typically three phases were observed during encounters: the initial phase, defined between the removal of partitions and the first aggressive display, the second phase or social conflict, in which symmetric aggressive behaviors were observed (lateral displays, circles, strikes, chases and bites), and the third phase starting with the conflict resolution, which is defined when a winner and a loser emerge as an asymmetry of expressed behaviors, with all aggressive acts initiated by the same fish (Oliveira et al., 2011). All behavioral interactions for 30 minutes were video-recorded with a JVC HD Everio camera for subsequent behavioral analysis.

For behavioral data, video recordings were analyzed by a blind observer to minimize observer bias and behavioral analysis was performed following the original ethogram for dyadic agonistic encounters in males of this species (Oliveira et al., 2011). The following behavioral variables were quantified during the agonistic encounters: (1) latency for the first attack (i.e., the time between the beginning of the recording period and the first aggressive behavior); (2) time of aggression (total time during which fish engage in any aggressive behavior, i.e., display, circle, bite, chase or strike); (3) time of freezing. Moreover, the following individual variables were quantified using Ethovision during 5 minutes before and after each treatment was applied: (4) distance travelled; (5) mean velocity (considering only the time frames in which each animal is in movement and excluding those with freezing); (6) freezing (speed lower than 0.2 cm/sec, Akinrinade et al., 2023a).

### 2.5. Specimen processing and tissue collection

Immediately after the second agonistic encounter, social interactions were stopped by placing an opaque partition in each tank and separating both opponents. Fish remained in isolation for 30 minutes after which they were euthanized by a rapid chilling followed by decapitation (American Veterinary Medical Association, AVMA, Guidelines for the Euthanasia of Animals, 2020). The choice of 30 minutes post-interaction was based on brain sampling, since brains will be used in further studies. Sex was corroborated by gonadal inspection and gonads were dissected, weighted and used for the calculation of gonadosomatic index (GSI%: gonad weight/body weight x 100). Morphometric variables were also determined in each fish (total and standard length and body weight). Female reproductive status was first determined by body shape (prespawning females are more ventrally rounded than postspawning females), and then by the GSI related to the reproductive status (Brewer et al., 2008, Cavallino et al., 2019). The mean values of GSI for prespawning and postspawning females was 12.76±0.39 mg and 6.77±0.52 mg, respectively.

### 2.6. Statistical Analyses

The GSI% was used to measure relative gonad size. Normality for behavioral data was assessed by Shapiro Wilks test, while homoscedasticity was evaluated graphically by analyzing residuals and by Levene test. Since data did not meet normality, to study possible differences in agonistic behavior, ‘Treatment’, ‘Day’ and ‘Reproductive status’ were used as factors in a generalized linear mixed model (GLMM) which took into account their interactions, and ‘Pair’ as a random factor. To study possible differences in individual behavior, and considering that delta values met normality, ‘Treatment’ and ‘Day’ were used as factors in a generalized linear mixed model (LMM) taking into account their interaction, and ‘Pair’ as aleatory variable. In all cases, a posteriori Tukey contrasts were used. All statistical analyses were performed using the computer program R (v 3.6.1), including latest versions of packages emmeans, glmmTMB, DHARMa, car, lme4.

## Results

### 2.7. Effect of alarm substances in agonistic behavior during consecutive agonistic encounters

Potential differences in agonistic behavior due to fear response were assessed by exposing both opponents to AS, and this was compared with a control with no influx (Ctrl 1) and with a group exposed to water influx (Ctrl 2). Also, to further determine which is the effect of AS on subsequent agonistic encounters, opponents were exposed to the same treatment during a second trial the following day. In this sense, it is worth mentioning that in the general analysis there are non-significant interactions between Day and Treatment for all variables (p=0.8497 for latency, p=0.0577 for time of aggression, p=0.4139 for freezing, Table 1). Results suggest that AS increased the latency to the first attack (p<0.0001 when compared to Ctrl 1 and to Ctrl 2, Fig. 2A, Table 1 and 2) and individual freezing (p=0.0124 and p=0.0011 when compared to Ctrl 1 and Ctr 2, respectively; Fig. 2B, Table 1 and 2). However, exposure to AS does not alter the total time of aggression (p= 0.9336, Fig. 2C, Table 1, Table 2).

**Figure 2.**
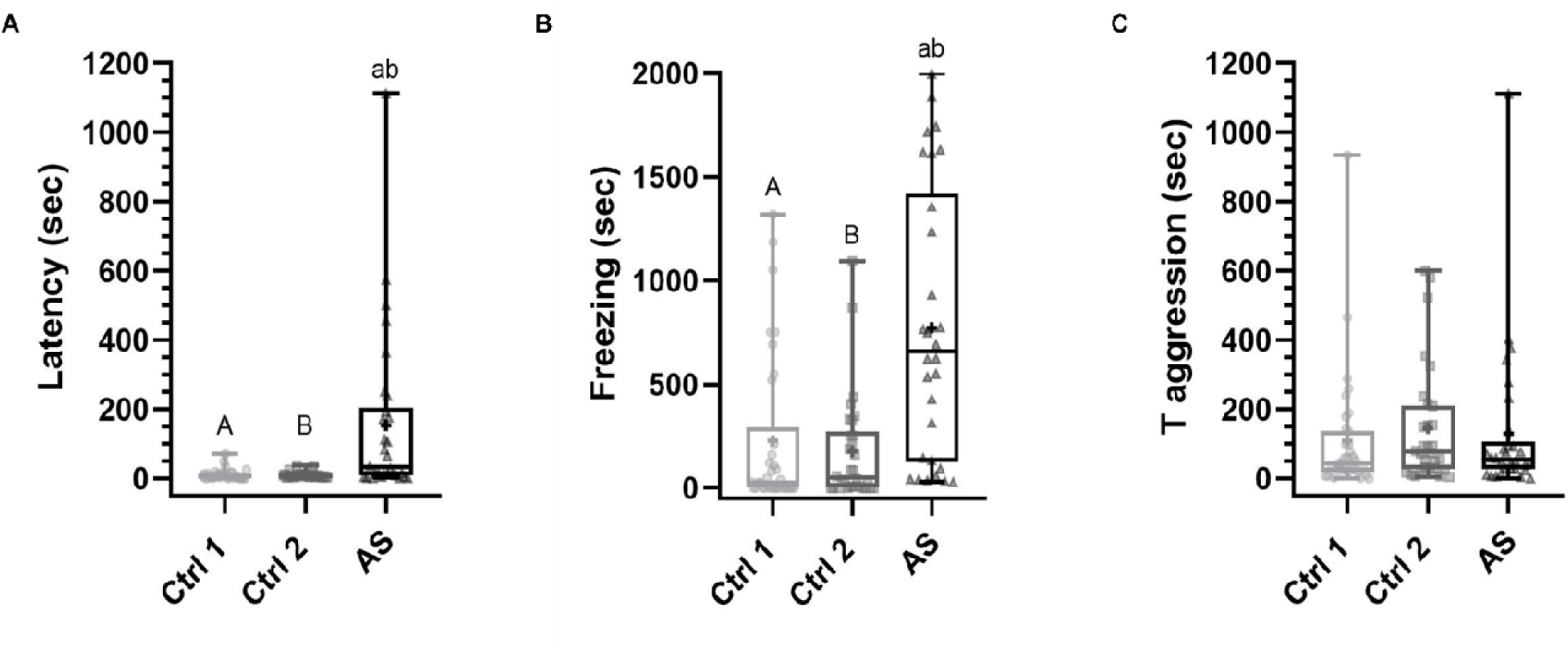
Effect of alarm substances during intrasexual dyadic agonistic encounters. A. Latency to the first attack. B. Freezing. C. Time of aggression. Treatments were administered via a flexible and transparent PVC tubing positioned 0.5 cm above the water level at each half of the tank. Ctrl 1: no added treatment. Ctrl 2: water influx. AS: alarm substances. Individual values are represented by dots (Ctrl 1), squares (Ctrl 2) or triangles (AS). Horizontal lines express median, crosses represent the mean value, boxes indicate interquartile range and whiskers the maximum and minimum values. Data was compared by GLMM followed by Tukey contrasts. Differe t letters show different contrasts. Capital and lowercase letters show statistical differences.

**Table 1.** Statistical analysis comparing agonistic behavior during consecutive agonistic encounters. A. GLMM analysis. B. A posteriori Tukey contrasts. Ctrl1: no added treatment. Ctrl2: water influx. AS: alarm substances. Sec: seconds. Post-post: dyads in which both opponents are in post spawning. Pre-post: dyads in which one female is in pre spawning and the other one is in post spawning. Pre-pre: dyads in which both opponents are in pre spawning.

|  | Latency (sec) |  | Time of aggression (sec) |  | Freezing (sec) |  |
| --- | --- | --- | --- | --- | --- | --- |
|  | Chi-square | p-value | Chi-square | p-value | Chi-square | p-value |
| 1A |  |  |  |  |  |  |
| Day | 11.3475 | 0.0008 | 7.6648 | 0.0056 | 0.0828 | 0.7735 |
| Treatment | 62.5078 | 2.67E-11 | 0.1374 | 0.9336 | 11.7786 | 0.0028 |
| Reproductive status | 7.8286 | 0.02 | 6.6738 | 0.0355 | 2.4713 | 0.2906 |
| Day*Treatment | 0.3256 | 0.8497 | 5.7741 | 0.0557 | 1.7642 | 0.4139 |
| Treatment*Reproductive status | 4.1672 | 0.3839 | 1.0702 | 0.899 | 9.1798 | 0.0568 |
| 1B | z test | p-value | z test | p-value | z test | p-value |
| Ctrl1 vs Ctrl2 | -0.229 | 0.9715 | - | - | -0.692 | 0.768 |
| Ctrl1 vs AS | 6.755 | <.0001 | - | - | 2.842 | 0.0124 |
| Ctrl2 vs AS | -5.28 | <.0001 | - | - | -3.566 | 0.0011 |
| Day 1 vs Day 2 | -3.28 | 0.001 | 2.631 | 0.0085 | - | - |
| post-post vs pre-post | -0.394 | 0.9178 | 2.55 | 0.029 | - | - |
| post-post vs pre-pre | 1.003 | 0.575 | 0.966 | 0.5984 | - | - |
| pre-post vs pre-pre | 1.232 | 0.4345 | -1.236 | 0.4316 | - | - |

**Table 2.** Values for the statistical analysis comparing agonistic behavior during consecutive agonistic encounters. Ctrl1: no added treatment. Ctrl2: water influx. AS: alarm substances. Sec: seconds. CI: confidence interval. IR: interquartile range.

|  |  | Treatment |  |  | Day |  |
| --- | --- | --- | --- | --- | --- | --- |
|  |  | Ctrl1 | Ctrl2 | AS | Day1 | Day2 |
| Latency (sec) | Mean | 11.59 | 10.5 | 146.7 | 43.53 | 69.3 |
|  | CI | (6.54, 16.64) | (7.25, 13.75) | (55.34, 238.06) | (12.0, 75.06) | (14.7, 123.89) |
|  | Median | 6.5 | 8 | 22.5 | 6 | 13 |
|  | IR | 10.5 | 6.5 | 168.25 | 8 | 28 |
|  | n | 34 | 30 | 30 | 47 | 47 |
| Time of aggression (sec) | Mean | 108.3 | 145.23 | 134.03 | 161.85 | 95.07 |
|  | CI | (44.18, 172.43) | (81.58, 208.88) | (50.47, 217.6) | (93.82, 229.87) | (55.48, 134.65) |
|  | Median | 43 | 76.5 | 59 | 73.5 | 49.5 |
|  | IR | 78 | 166.25 | 66 | 198.5 | 66.25 |
|  | n | 33 | 30 | 29 | 46 | 46 |
| Freezing (sec) | Mean | 228.25 | 175.33 | 772.58 | 334.26 | 435.91 |
|  | CI | (95.46, 361.04) | (77.92, 272.75) | (529.34, 1015.82) | (204.95, 463.57) | (259.78, 612.05) |
|  | Median | 26 | 50.75 | 658.5 | 86 | 110 |
|  | IR | 199.5 | 249.63 | 1192.25 | 549.75 | 732.25 |
|  | n | 34 | 30 | 30 | 47 | 47 |

Results also suggest that there is a significant effect of the Day in both latency (p=0.0008, Table 1) and time of aggression (p=0.0056, Table 1), but not in freezing (p=0.7735, Table 1). This way, a second exposure to an agonistic encounter in all treatments increases both latency to the first attack and reduces the time invested in aggressive displays (Table 2, Fig. 3).

**Figure 3.**
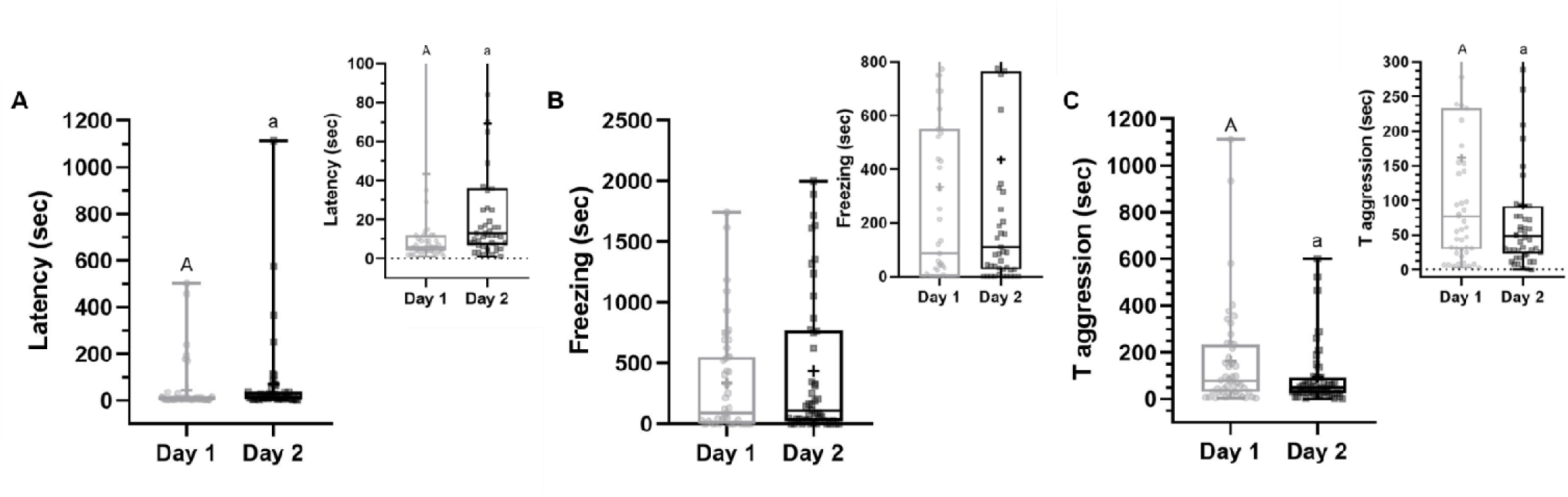
Agonistic behavior during consecutive intrasexual dyadic agonistic encounters. A. Latency to the first attack. B. Freezing. C. Time of aggression. Individual values are represented by dots (Day 1) or squares (Day 2). Horizontal lines express median, crosses represent the mean value, boxes indicate interquartile range and whiskers the maximum and minimum values. Data was compared by GLMM followed by Tukey contrasts. Capital and lowercase letters show statistical differences.

To assess whether agonistic behavior is altered according to the different reproductive status of both opponents, this was included as a factor in the analysis. While the general analysis suggests that there are no significant differences in freezing (p=0.2906), it shows significant differences in latency according to the reproductive status (p=0.02) which is not detected by *a posteriori* contrasts (Table 1). Results also suggest that there are significant differences in time of aggression (p=0.0355), in particular when comparing dyads in which both opponents correspond to the post spawning stage and mixed dyads with both females in different reproductive stages (p=0.029, Table 1). However, there are non-significant interactions between Treatment and Reproductive status for all variables (p=0.3839 for latency, p=0.899 for time of aggression, p=0.0568 for freezing). Overall, these results suggest that exposure to AS alters agonistic behavior in females by increasing both freezing and latency to the first attack, and that a second encounter increases latency and reduces time of aggression.

### 2.8. Effect of alarm substances in individual behavior before consecutive agonistic encounters

To assess whether the effect of fear response on individual behavior is due to the alarm substances themselves or to the treatment influx, individual distance travelled, mean velocity and freezing were compared between before and after the stimuli, before performing the agonistic encounter. Results suggest that both distance travelled and mean velocity are significantly reduced after adding AS (p = 0.00006, p= 0.0419, respectively), but not because of the water influx (p=0.1591, p= 0.4037, respectively, Fig. 4A,B,D,E, Table 3). Interestingly, freezing is increased after treatment administration in both water control and AS (p=0.0214, p=0.0002, respectively, Fig 4C,F, Table 3).

**Figure 4.**
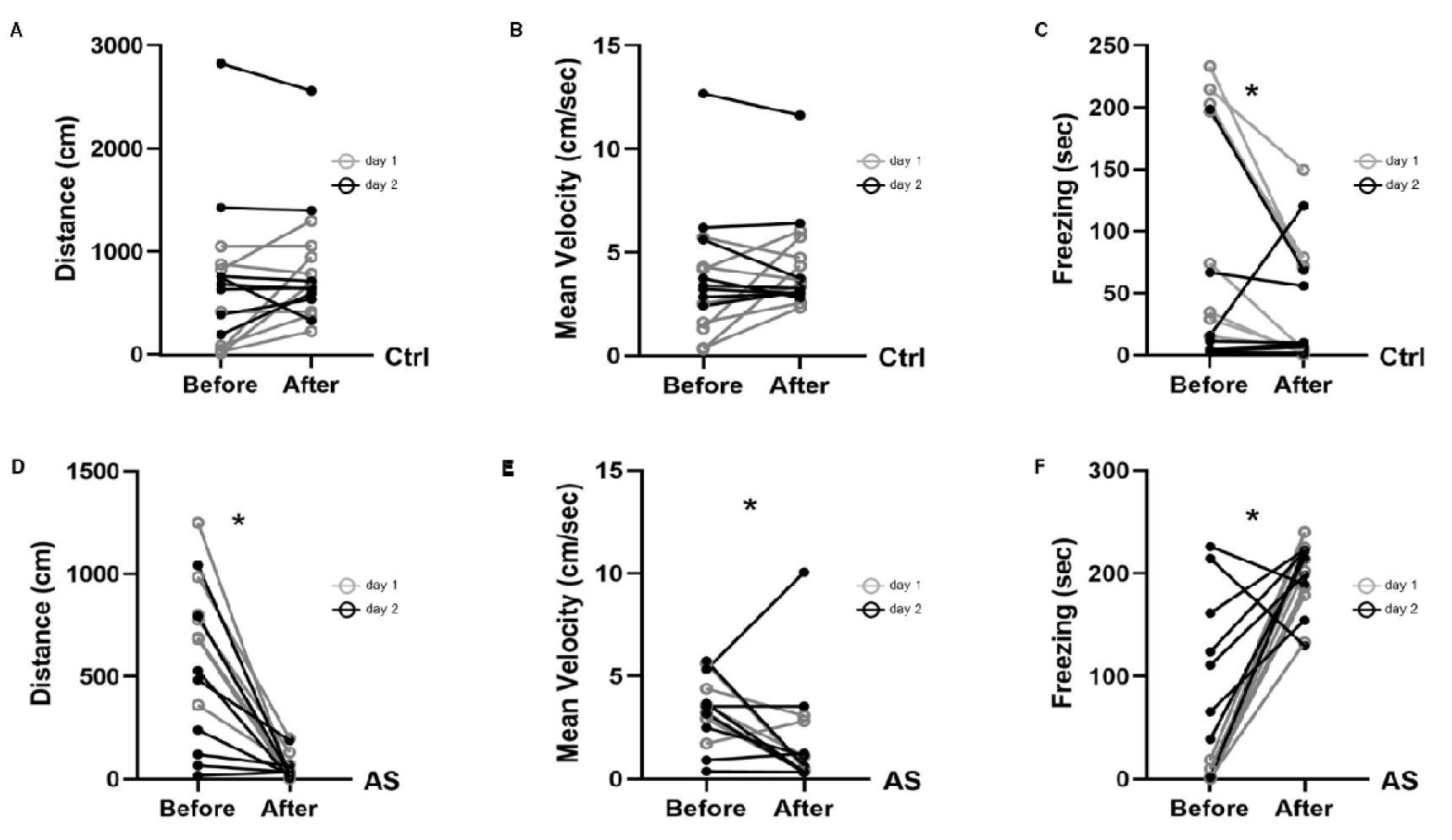
Effect of alarm substances on individual behavior. Distance, mean velocity and freezing were determined during five minutes before and after stimuli administration. A, B, C. Both opponents were exposed to water addition with no alarm substances. D, R, F. Both opponents were exposed to alarm substances (AS). Individual values of each fish are plotted as dots connected by a line. Data was compared with paired Wilcoxon test and asterisks indicate statistical differences.

**Table 3.**
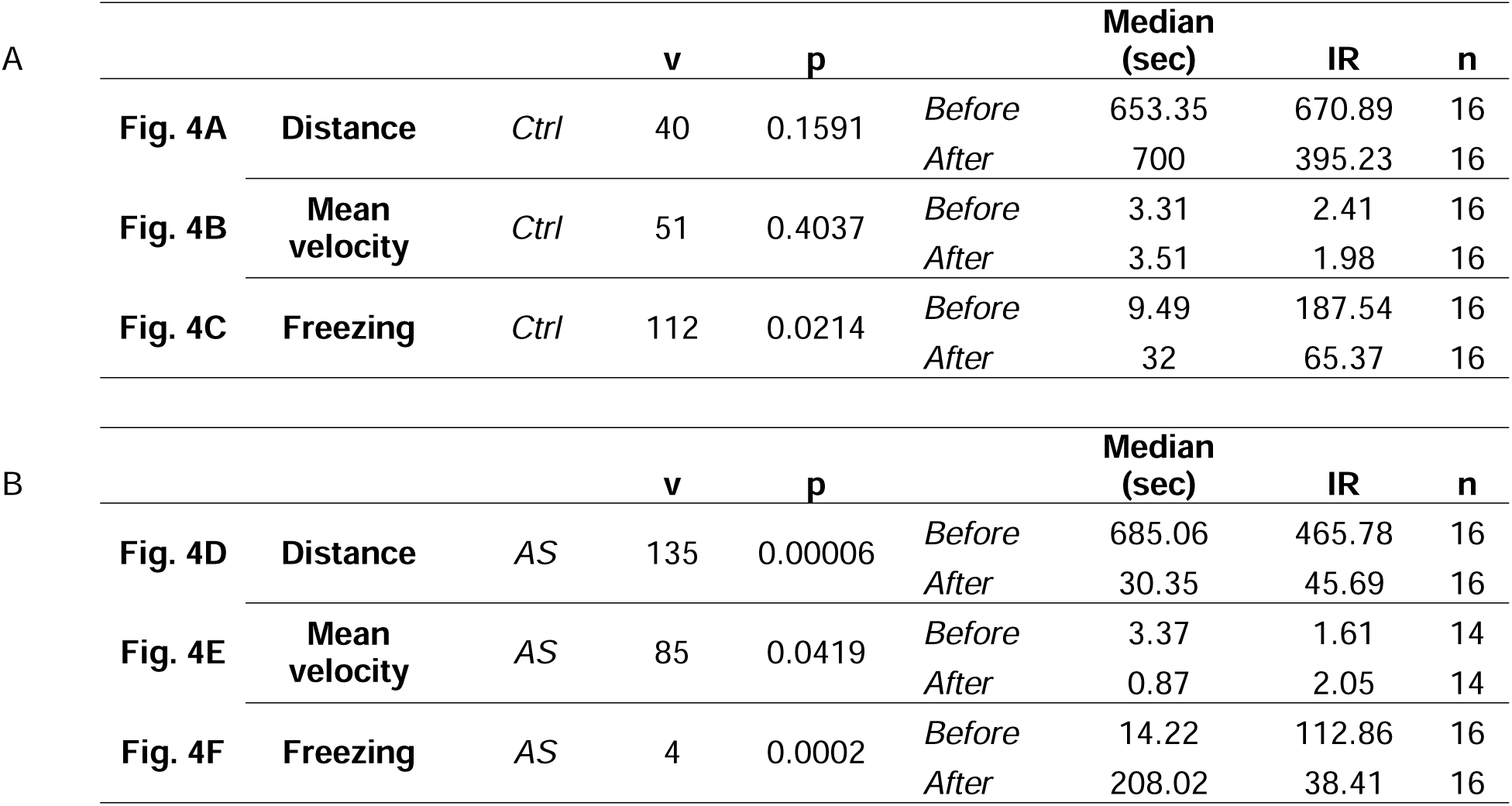
Paired Wilcoxon analysis comparing individual behavior during five minutes before and after treatment. V and p-values for each behavioral display before and after administration of control water (A, Fig. 4A-C) or AS (B, Fig. 4D-F).

Moreover, changes in behavior before and after adding stimuli are expressed with delta values (Δ), calculated as each variable post stimuli minus values pre stimuli. This was assessed before performing the agonistic encounter, both in day 1 and day 2. It is worth emphasizing that in the general analysis there are significant interactions between Day and Treatment for all variables (p=0.0005 for delta distance, p=0.0145 for delta mean velocity, p= 0.000029 for delta freezing, Table 3). When comparing the changes in distance before and after adding stimuli, results suggest significant differences between fish exposed to water influx and AS in the first day (p= <0.0001) and in the second day (p=0.04, Table 4 and 5, Fig. 5A). Moreover, changes in distance before and after adding the water influx in the second day differ significantly when compared to the first day (p= 0.0246, Table 4 and 5, Fig. 5A). Similar results are observed after adding AS in the second day, when compared to the first day (p= 0.031, Table 4 and 5, Fig. 5A). Regarding changes in mean velocity, results suggest significant differences between treatments in the first day (p= 0.0004, Table 4 and 5, Fig. 5B). When comparing the change in freezing before and after adding stimuli, results suggest significant differences between treatments in the first day (p <0.0001, Table 4 and 5, Fig. 5C) and also in the second day (p= 0.0252, Table 4 and 5, Fig. 5C). Results also suggest that changes in freezing before and after adding the water influx in the second day differ significantly when compared to the first day (p= 0.0433, Table 4 and 5, Fig. 5C). Similar results are observed after adding AS in the second day, when compared to the first day (p=0.0024, Table 4 and 5, Fig. 5C).

**Table 4.**
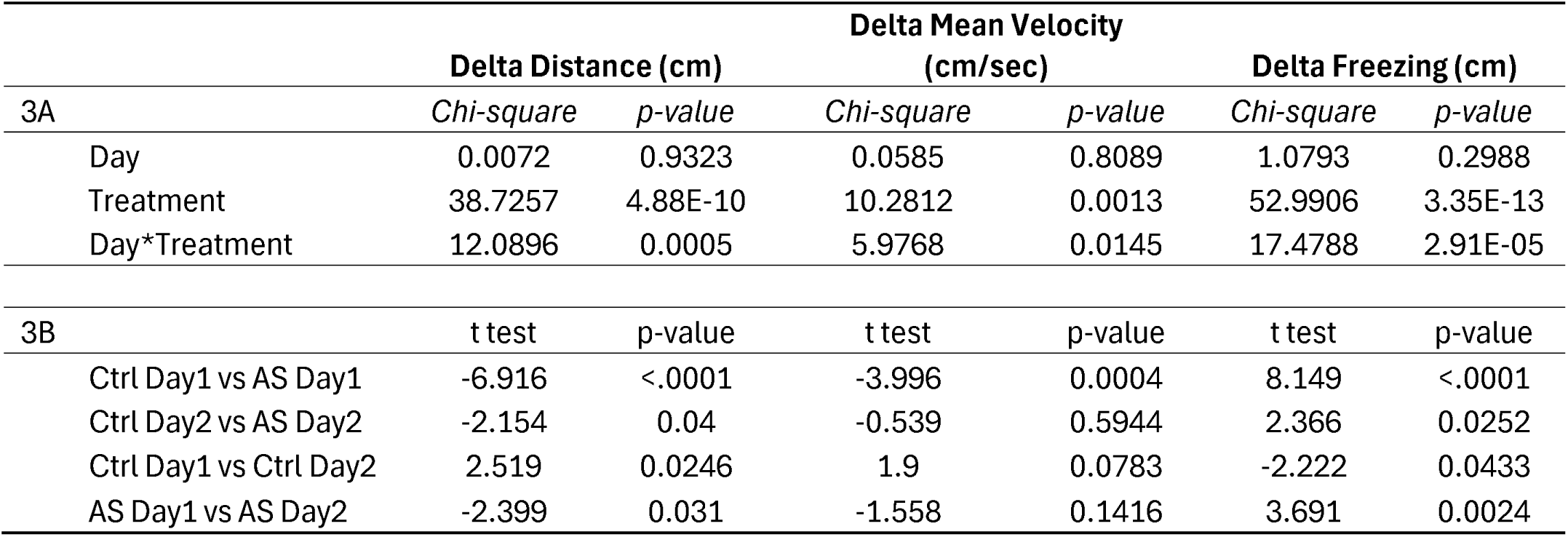
Statistical analysis comparing individual behavior during consecutive agonistic encounters. Changes in each behavior are expressed with delta, calculated as each variable post stimuli minus values pre stimuli. A. GLMM analysis. B. A posteriori Tukey contrasts. Ctrl: water influx. AS: alarm substances. Sec: seconds.

| 3A | Delta Distance (cm) |  | Delta Mean Velocity (cm/sec) |  | Delta Freezing (cm) |  |
| --- | --- | --- | --- | --- | --- | --- |
|  | Chi-square | p-value | Chi-square | p-value | Chi-square | p-value |
| Day | 0.0072 | 0.9323 | 0.0585 | 0.8089 | 1.0793 | 0.2988 |
| Treatment | 38.7257 | 4.88E-10 | 10.2812 | 0.0013 | 52.9906 | 3.35E-13 |
| Day*Treatment | 12.0896 | 0.0005 | 5.9768 | 0.0145 | 17.4788 | 2.91E-05 |

*Table 4. Statistical analysis comparing individual behavior during consecutive agonistic encounters.
| 3B | t test | p-value | t test | p-value | t test | p-value |
| --- | --- | --- | --- | --- | --- | --- |
| Ctrl Day1 vs AS Day1 | -6.916 | <.0001 | -3.996 | 0.0004 | 8.149 | <.0001 |
| Ctrl Day2 vs AS Day2 | -2.154 | 0.04 | -0.539 | 0.5944 | 2.366 | 0.0252 |
| Ctrl Day1 vs Ctrl Day2 | 2.519 | 0.0246 | 1.9 | 0.0783 | -2.222 | 0.0433 |
| AS Day1 vs AS Day2 | -2.399 | 0.031 | -1.558 | 0.1416 | 3.691 | 0.0024 |

**Table 5.** Values for the statistical analysis comparing individual behavior during consecutive agonistic encounters. Ctrl: water influx. AS: alarm substances. Sec: seconds. CI: confidence interval. IR: interquartile range.

|  |  | Ctrl Day1 | AS Day1 | Ctrl Day2 | AS Day2 |
| --- | --- | --- | --- | --- | --- |
| **Delta Distance (cm)** | *Mean* | 345.76 | -719.03 | -29.76 | -361.42 |
|  | *CI* | (68.4, 623.12) | (-935.51, -502.54) | (-235.71, 176.2) | (-675.84, -47.0) |
|  | *Median* | 300.34 | -673.6 | -36.16 | -255.78 |
|  | *IR* | 374.44 | 140.78 | 149.11 | 533.49 |
|  | *n* | 8 | 8 | 8 | 8 |
| **Delta Mean Velocity (cm/sec)** | *Mean* | 1.55 | -2.55 | -0.4 | -0.95 |
|  | *CI* | (-0.09, 3.19) | (-4.09, -1.0) | (-1.09, 0.3) | (-3.44, 1.54) |
|  | *Median* | 1.39 | -2.69 | -0.16 | -0.69 |
|  | *IR* | 2 | 1.22 | 1.12 | 3.05 |
|  | *n* | 8 | 8 | 8 | 8 |
| **Delta Freezing (cm)** | *Mean* | -76.83 | 198.06 | -3.52 | 76.28 |
|  | *CI* | (-122.74, -30.92) | (168.91, 226.22) | (-56.27, 49.23) | (-7.64, 160.2) |
|  | *Median* | -67.48 | 202.23 | 1.05 | 88.2 |
|  | *IR* | 89.32 | 38.74 | 6.94 | 80.03 |
|  | *n* | 8 | 8 | 8 | 8 |

**Figure 5.**
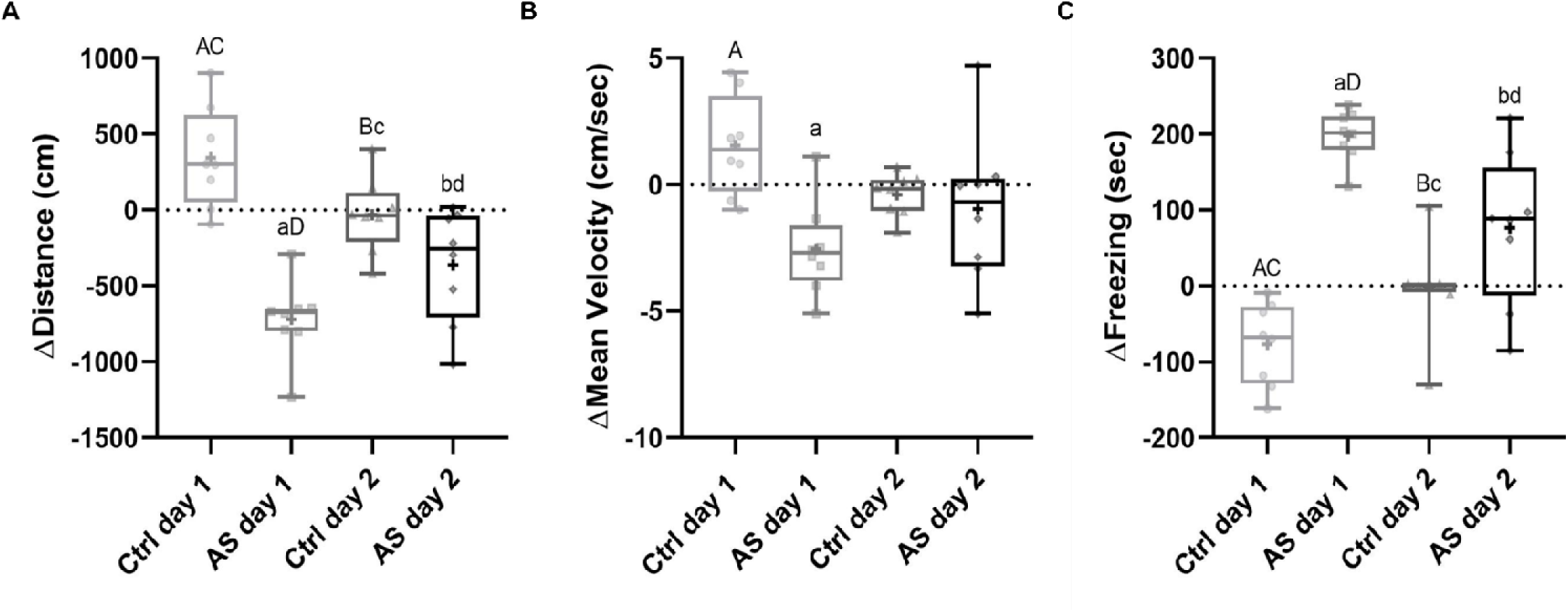
Changes in behavior due to consecutive exposure to alarm substances. Changes are expressed with delta (Δ), calculated as each variable post stimuli minus values pre stimuli. Ctrl refers to dyads with both opponents exposed to water addition with no alarm substances. AS refers to dyads with both opponents exposed to alarm substances. Individual values are represented by dots (Ctrl day1), squares (AS day 1), triangles (Ctrl day 2) and diamonds (AS day 2). Horizontal lines express median, crosses represent the mean value, boxes indicate interquartile range and whiskers the maximum and minimum values. Data was compared by GLMM followed by Tukey contrasts. Different letters show different contrasts. Capital and lowercase letters show statistical differences.

## 3. Discussion

We assessed how the repeated exposure to the distress caused by fear sensing (e.g. exposure to alarm substances) alters different aspects of individual and aggressive behavior during consecutive agonistic encounters in female zebrafish. Results in this study suggest that alarm substances reduce female motivation to engage in an agonistic encounter, and that the aggressive behavior is still maintained despite sensing alarm substances as potential threat. This effect of the alarm substance is observed for the first day and for the second day of agonistic encounter. Finally, similar to agonistic behavior, the effect of alarm substance on certain individual behavioral parameters such as distance and freezing is also observed during a first and also a second exposure to alarm substances.

The fact that females in the prespawning and in the postspawning stage show no differences in the latency and in the time of aggression suggests a similar motivation to engage in an agonistic encounter and similar levels of aggression regardless of the reproductive stage. This is aligned with evidence in other teleost species such as the Neotropical cichlid *C. dimerus*, in which it has been suggested that certain parameters indicating the reproductive stage in females may be not very informative to explain territorial aggressive behavior in a dyadic neutral context (Scaia et al., 2023). Interestingly, the relation between aggression and the reproductive stage seems to depend on the ethological approach and on the species (Tubert et al., 2012, Seebacher et al., 2013). Even though historically studies on aggressive behavior usually tend to focus on males, in zebrafish there is growing evidence on the neuroendocrine and neural basis to understand female aggression (Filby et al., 2010, Paull et al., 2010, Umeda et al., 2022, Scaia et al., 2022). In this context, it is important to recall that one of the most evident morphological differences between sexes in zebrafish besides body color is the fact that females typically show a marked distension and more rounded abdomen than males, who present slender body shape (Crowder et al., 2018). Considering that prespwning females show more developed gonads and thus a more prominent distension of their abdomen than postspawning females, evidence in female zebrafish tends to focus on the prespawning stage. To the best of our knowledge, this is the first study comparing agonistic behavior between both reproductive status in zebrafish females. Interestingly, time of aggression differs when comparing encounters with both females in postspawning stage, and those with one in prespawning and its opponent in postspawning. This evidence could suggest that levels of female aggression might differ when sensing an opponent in a different reproductive stage than the focal individual, but this should be assessed in future studies.

A very popular paradigm to study fear response in fish is the exposure to alarm substances, which induce erratic movements and freezing (Speedie and Gerlai, 2008). Since this alarm reaction is related to an increase in cortisol levels and stress (Egan et al., 2009), the use of alarm substances in zebrafish is a very popular approach to study distress behavior and fear sensing. Fear response, referring to the behavioral modulation in presence of a potential risk in the environment, is a very appealing framework to assess different manifestations of social information use such as social buffering, social transmission (contagion) and facilitation (Oliveira and Faustino, 2017). When assessing fear response induced by AS, behavioral studies traditionally focus on the effect of alarm substances on individual traits, suggesting an increased freezing, erratic movements and a reduction in time spent in the upper half of the tank (Egan et al., 2009, Speedie and Gerlai, 2008). Besides individual behavior, different aspects of social behavior have been assessed in the context of distress and fear sensing in fish. In this scenario, the presence of a conspecific in a social context may act either as a threat ameliorating stimulus or as a stressor itself (Graves and Hennessy, 2000). When a potential threat is presented to a focal animal simultaneously to the presence of conspecifics, focal individuals combine direct and indirect or social information to assess the presence of a potential threat in the environment and modulate their behavior accordingly, resulting in different social phenomena. While social buffering refers to the reduction of threat response when sensing the relaxed behavior of conspecifics, social contagion occurs when the alarm behavior of conspecifics contradicts the lack of threat detected by the individual, thus triggering an alarm response even in the absence of a direct threat (Oliveira and Faustino, 2017).

Evidence on the neural mechanisms underlying these social phenomena in zebrafish suggest a shared evolutionary origin for social buffering, and also for the role of oxytocin as one of the main regulators to understand social fear contagion (Faustino et al., 2017, Akinrinade et al., 2023a,b). This growing literature constitutes a promising framework to study to which extent agonistic behavior can be altered by distress using zebrafish as a biological model. Results in this study suggest that distress caused by the exposure to alarm substances increases the latency in dyadic contests, suggesting that sensing chemical cues related to a potential threat reduces the motivation to engage in an agonistic encounter. This can be understood in the context of integrating relative costs and benefit trade-offs between sensing a potential danger and engaging in a contest (Hsu et al., 2021, Brown et al., 2006, Wishingrad et al., 2014). Interestingly, this effect of AS does not differ between the first and the second exposure to the treatment and to agonistic encounter. However, since aggression is not altered in presence of AS, this suggests that despite the fact that fish sense danger or an aversive potential risk in the environment, they still engage in fights and dedicate time to aggressive encounters. It seems that the threat of predation (e.g chemical cues from injured conspecifics) temporarily delays the decision to engage in a contest thus reducing initial motivation, but once fish observe no real injured conspecific but a real opponent instead, they decide to engage in agonistic encounters.

The fact that time of aggression is not altered in presence of alarm substances can also have an alternative interpretation, since the presence of another female in the same tank can represent not only an opponent itself but it may also have an effect as social buffer. However, results suggest that freezing is significantly increased in the presence of AS, suggesting that the presence of a conspecific does not reduce distress and fear response. Available literature referring to aggressive behavior during fear sensing is scarce. Evidence in zebrafish suggests that aggression is reduced when alarm substances are added to the zebrafish shoal but not when there are only two opponents (Rehnberg and Smith, 1988). This is similar to results in this paper, even if here we include not only chases but all aggressive displays described for this species (Oliveira et al., 2011) allowing us to determine the significant effect of AS in latency not only during the first, but also during the second encounter. Other evidence in zebrafish suggests that an acute exposure to AS increases aggression and does not affect distance travelled and latency to the first attack, while chronic exposure to AS reduced aggression and locomotion (Quadros et al., 2018). This apparent discrepancy in the effect of exposure to AS on aggression can be better understood considering that evidence refers to mirror fights while in this study we assess dyadic encounters with real opponents. As such, exposure to AS can have a differential effect on agonistic behavior if the opponent is a real or a virtual conspecific, these differences probably explained by the presence of other chemical cues in the case of a real opponent, or by the fact that a real opponent does not mirror the behavior of a focal fish.

There is also evidence in convict cichlid fish suggesting that species-specific antipredator response to conspecific alarm substances reduces intraspecific aggression (Wisenden and Sargent, 1997). Interestingly, besides the phylogenetic distance, in this case authors study juveniles and instead of comparing total time of aggression among treatments, they compare the change in approaches, bites and chases before and after exposing fish to the stimuli. Taking these methodological and ethological differences into account with previous evidence, results in the present study suggest that alarm substances reduce female motivation to engage in an agonistic encounter, and that the aggressive behavior is still maintained despite sensing a potential threat.

In general fear response in fish is assessed by acute exposure to alarm substances, since chronic exposure for 30 minutes has been suggested to be less effective inducing freezing and erratic movements (Egan et al., 2009). Results here show that the alarm substances increase freezing when considering long exposures throughout the duration of encounters, and this increase in freezing is also evident during the second day of contests. In this regard it is important to recall that during the second day fish were exposed not only to the fresh AS but also to the AS used for the previous encounter, thus a possible cumulative effect cannot be excluded. Results comparing five minutes before and after adding the alarm substances and during physical isolation from the opponent also suggest an increase in freezing. Even if this effect cannot be excluded to be at least in part due to the volume influx, change in freezing before and after the stimuli is higher in fish exposed to AS in the first and also in the second day. These differences in the increase in freezing during these experiments and the reported evidence on the chronic exposure can also be argued considering that authors originally considered freezing as total absence of movement for at least 1 sec, while here we use more recent evidence considering freezing as when velocity is less than 0.2 cm/sec.

Regarding consecutive agonistic encounters, evidence on zebrafish males show a reduction in time of aggression during a second agonistic encounter, suggesting the formation of a long-term memory related to recognizing a particular opponent or features of a previous fighting experience (Cavallino et al., 2020, 2024). In this context, the present study represents the first evidence assessing how consecutive agonistic encounters with a real opponent can be altered by repeated exposure to alarm substances. Interestingly, even if the effect of AS on agonistic behavior is observed for the first and the second day of exposure to the treatment and to social interactions with no differences between days, results in this study suggest that in the second day there is a reduction in the time of aggression regardless of treatments. This reduction in aggression during a second agonistic encounter is similar to what has been reported for males of this species (Cavallino et al., 2020). Results here also show that in females there is a reduction in motivation to initiate the first agonistic encounter during the second day regardless of the treatment. However, since in males there are no differences in latency, this discrepancy could be related not only to the sex but also to the different statistical analysis used, since in that case latency was compared during three days of agonistic encounters.

Finally, when analyzing individual behavioral data before and after adding alarm substances during physical isolation of both opponents, results suggest that AS alter the change in distance and freezing during both the first and the second exposure, while the change in mean velocity is only altered during the exposure to AS only the first day. The fact that during the second exposure there are differences in the effect of AS on freezing and distance but not on velocity might be related to the fact that velocity reflects the mean value, while freezing and distance refer to the overall values of the time set. Overall, results in this study suggest that the effect the exposure to AS on agonistic behavior and individual behavioral traits is maintained during a second exposure to alarm substances. This suggests there is no attenuation of the effect of alarm substances causing distress and that alarm substances do not lose physiological effect on fear sensing in both opponents during social interactions.

## Acknowledgements

This work was supported by the Agencia de Promoción Científica y Tecnológica (PICT 2021- 00043), CONICET (PIBAA 28720210100710CO), Universidad de Buenos Aires (UBACyT 20020220400073BA) and International Brain Research Organization (IBRO) Early Career.

## List of symbols and abbreviations

AS: alarm substances
GSI: gonadosomatic index
Ctrl: control
Δ: delta
Pre-Pre: prespawning – prespawning
Post-Post: postspawning – postspawning
T aggression: time of aggression.

## Notes

Ethical statement: Authors declare no conflict of interest.

### Competing Interest Statement

The authors have declared no competing interest.

